# Pat1 increases the range of decay factors and RNA bound by the Lsm1-7 complex

**DOI:** 10.1101/2020.04.07.029900

**Authors:** Joseph H. Lobel, John D. Gross

## Abstract

Pat1 promotes the activation and assembly of multiple proteins during mRNA decay. After deadenylation, the Pat1/Lsm1-7 complex binds to transcripts containing oligo(A) tails, which can be modified by the addition of several terminal uridine residues. Pat1 enhances Lsm1-7 binding to the 3’ end, but it is unknown how this interaction is influenced by nucleotide composition. Here we examine Pat1/Lsm1-7 binding to a series of oligoribonucleotides containing different A/U contents using recombinant purified proteins from fission yeast. We observe a positive correlation between fractional uridine content and Lsm1-7 binding affinity. Addition of Pat1 broadens RNA specificity of Lsm1-7 by enhancing binding to A-rich RNAs and increases cooperativity on all oligonucleotides tested. Consistent with increased cooperativity, Pat1 promotes multimerization of the Lsm1-7 complex, which is potentiated by RNA binding. Furthermore, the inherent ability of Pat1 to multimerize drives liquid-liquid phase separation with multivalent decapping enzyme complexes of Dcp1/Dcp2. Our results uncover how Pat1 regulates RNA binding and higher order assembly by mRNA decay factors.

## INTRODUCTION

A dense network of protein-protein interactions regulates 5’-3’ mRNA decay, which is important for gene expression and many physiological processes including development, microRNA-mediated decay, and quality control mechanisms (Moore 2005; Kurosaki et al. 2019; Mugridge et al. 2018; Jonas and Izaurralde 2015). Bulk 5’-3’ mRNA degradation begins with the trimming of the 3’ poly(A) tail by cytoplasmic deadenylases, which can be followed by addition of several uridines by terminal uridine transferases in fission yeast and metazoans (Mugridge et al. 2018; Rissland and Norbury 2009; Lim et al. 2014; Yi et al. 2018; Webster et al. 2018). After deadenylation and subsequent uridylation, the heterooctameric Pat1/Lsm1-7 complex assembles on or near the 3’ A/U-rich deadenylated tail of the mRNA (Mitchell et al. 2012; Tharun and Parker 2001; Bonnerot et al. 2000; Bouveret et al. 2000; Tharun et al. 2000; Wang et al. 2017). Pat1 subsequently activates decapping by Dcp1/Dcp2, leading to rapid 5’-3’ degradation of the mRNA body by the conserved exonuclease Xrn1 (Lobel et al. 2019; Nissan et al. 2010; Stevens 1980; Mugridge et al. 2018; Charenton et al. 2017; Tharun and Parker 2001). Pat1 activates proteins at both the 5’ and 3’ end of the mRNA by enhancing RNA binding of the Lsm1-7 complex to the deadenylated 3’ end and decapping by the Dcp1/Dcp2 complex (Lobel et al. 2019; Nissan et al. 2010; Chowdhury et al. 2013; Charenton et al. 2017). Deletion of Pat1 results in accumulation of poorly translated, deadenylated, capped transcripts, suggesting a block in decapping (Wang et al. 2017; He et al. 2018; Tharun and Parker 2001).

Many mRNA decay factors, including Pat1, are enriched in Processing-bodies (P-bodies) which are a class of membraneless organelles that may function in mRNA storage or decay (Teixeira and Parker 2007; Xing et al. 2018; Hubstenberger et al. 2017; Sheth and Parker 2003). At the molecular level, these structures are promoted by multivalent protein-protein and protein-nucleic acid interactions that are required for phase separation (Banani et al. 2017). Overexpression of Pat1 enhances P-body formation in fungi, suggesting its importance in assembling these structures (Wang et al. 2017; Sachdev et al. 2019). Therefore, Pat1 functions at multiple steps during 5’-3’ mRNA decay to coordinate degradation of the transcript.

Pat1 uses a combination of disordered and globular domains to interact with and activate multiple mRNA decay factors. The disordered N-terminus contains a conserved FDF motif that interacts with Dhh1 (DDX6 in humans) and potentiates P-body formation, but is largely dispensable for function (Sachdev et al. 2019; Sharif et al. 2013; Pilkington and Parker 2008). The unstructured middle domain contains multiple short linear interaction motifs (SLiMs) and cooperates with the structured C-terminal domain to activate RNA binding by Lsm1-7 and decapping by the Dcp1/Dcp2 complex through multiple mechanisms (Lobel et al. 2019; Pilkington and Parker 2008; Chowdhury et al. 2013).

While it is known how Pat1 activates different mRNA decay factors, much less is understood about how it affects specific RNA recognition by Lsm1-7. *In vitro*, budding yeast Pat1/Lsm1-7 shows a preference for oligoadenyated RNAs compared to those containing poly(A) tails; however, genome-wide CLIP studies indicate Pat1/Lsm1 co-occupy the 3’ untranslated region (UTR) of budding yeast transcripts without enriching a specific sequence motif (Chowdhury et al. 2007; Mitchell et al. 2012). In fission yeast and metazoan cells, deletion of Lsm1 or Pat1 stabilizes mRNA decay intermediates with oligo(A) tails containing several uridines (Lim et al. 2014; Rissland and Norbury 2009). Furthermore, in mammalian cells, knockdown of Pat1 stabilizes transcripts with AU-rich sequences in the 3’ UTR (Vindry et al. 2017). Lsm1-7 can also bind oligo(U) RNA sequences *in vitro* and promotes decay of histone mRNAs that containing U-rich tails in cells (Mullen and Marzluff 2008; Chowdhury et al. 2007; Wu et al. 2014). How Pat1 affects location and sequence specificity of the Lsm1-7 complex on mRNA is poorly understood.

In this work, we evaluate recombinant purified *S.pombe* Pat1/Lsm1-7 complex binding to a series of oligonucleotides of different A/U content. Lsm1-7 alone has a binding preference for U-rich RNAs. Addition of Pat1, however, broadens the specificity of the Lsm1-7 complex by enhancing binding to A-rich targets. Furthermore, Pat1 increases cooperative binding of Lsm1-7 to oligonucleotides, which drives multimerization of the heterooctamer on RNA in a sequence independent manner. Oligomerization is an inherent property of Pat1 that permits higher order assembly with multivalent Dcp1/Dcp2 complexes, which can recruit additional mRNA decay machinery. Taken together, this work reveals how Pat1 broadens the specificity of Lsm1-7 and promotes the assembly of higher order decapping complexes.

## RESULTS

### The PatMC/Lsm1-7 complex cooperatively binds to A-rich RNA

Previous studies indicate that the middle and C-terminal domains of Pat1 (termed PatMC) are sufficient to support cell growth in yeast (Pilkington and Parker 2008; Lobel et al. 2019). To understand how different 3’ end sequences influence PatMC/Lsm1-7 binding, we tested recombinant purified *S.pombe* Lsm1-7 complexes alone or coexpressed with PatMC for their ability to bind different oligo-RNAs by fluorescence polarization (**Fig. 1A-B)**. Because global profiling of RNA tails indicate that uridine residues are found on short tails (<25 nt), we investigated a series of 15mers containing different adenine and uracil contents (Chang et al. 2014; Rissland and Norbury 2009). As seen previously, PatMC enhanced the RNA binding of Lsm1-7 to A15 RNA by an order of magnitude (Lobel et al. 2019). The fold-enhancement of Lsm1-7 RNA binding by PatMC strongly correlated with the fractional adenine content of the 15mer, where a greater difference in free energy of binding was observed for more adenine-rich substrates **(Fig. 1C & S1A-F)**. Furthermore, the Lsm1-7 complex alone strongly favored binding to U-rich 15mers, which was not affected by PatMC. Because PatMC does not bind RNA with appreciable affinity on its own, PatMC may serve to selectively enhance RNA binding of Lsm1-7 to adenine rich tails and may be dispensable for engaging U-rich tails (Lobel et al. 2019) (**Fig. S1G**).

**Figure 1:**
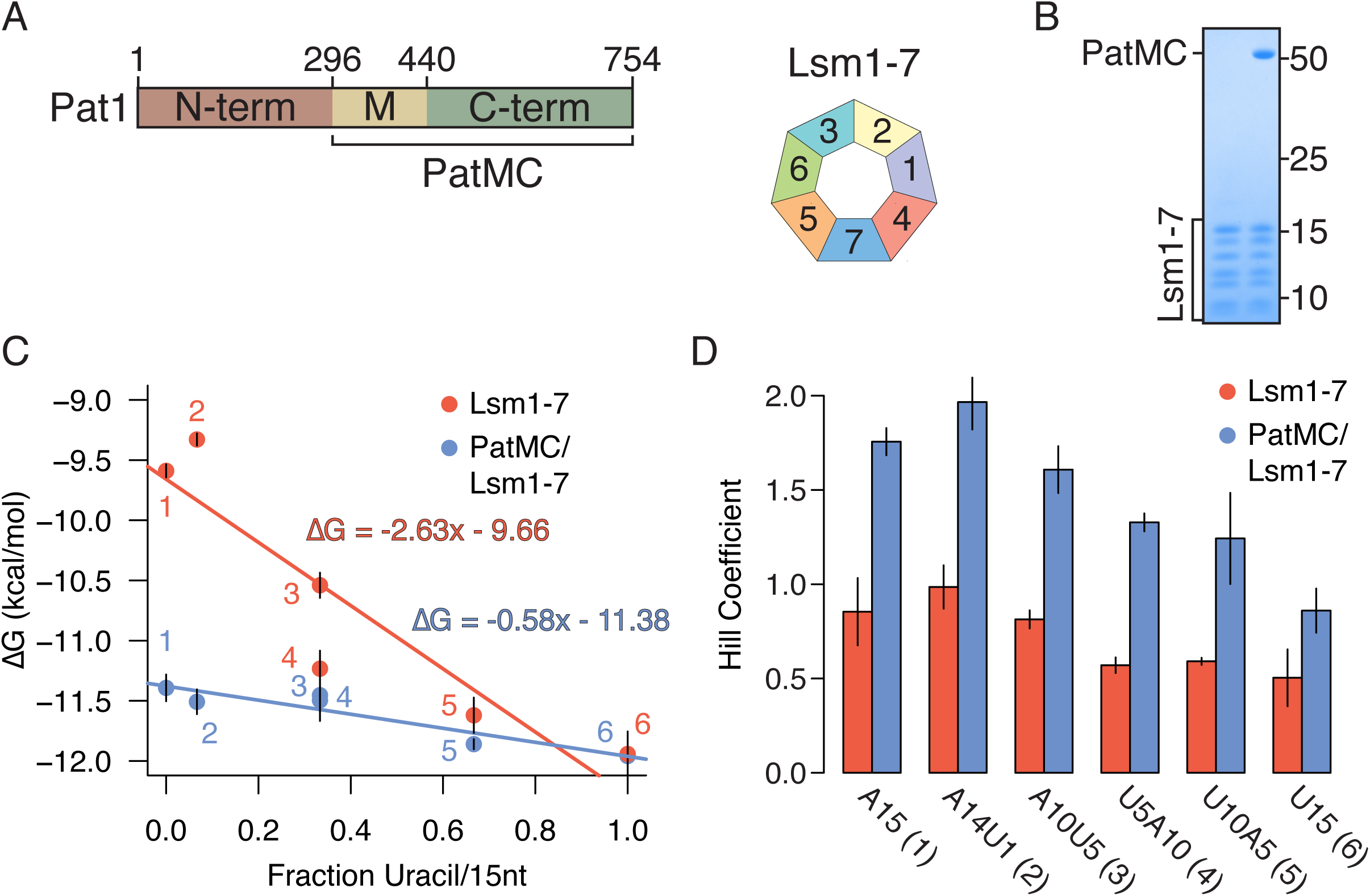
PatMC enhances Lsm1-7 binding to Adenine-rich substrates in a cooperative manner. **A**, Schematic of Pat1 domains and Lsm1-7 **B**, SDS-PAGE of the Lsm1-7 complex alone (left) or with PatMC (right). Molecular weight (kDa) shown on right. **C**, Lsm1-7 or PatMC/Lsm1-7 binding to different 5’-FAM labelled 15mer RNAs (A15, A14U1, A10U5, U5A10, U10A5, U15) monitored by fluorescence polarization. Numbers correspond to labels in **D**. Binding affinities were converted to ΔG and plotted against the fractional uracil content of the oligonucleotides (n=3). **D**, Hill coefficients for each fit for the binding isotherms shown in **C** (n=3).

In addition to the differences in affinities for the oligonucleotides, we also observed a difference in Hill coefficients, a measure of cooperativity that places a lower bound on the number of Pat1/Lsm1-7 complexes binding RNA. The binding isotherms of the PatMC/Lsm1-7 complex were consistently ∼2 fold more cooperative than Lsm1-7 alone for all tested RNAs, suggesting that PatMC/Lsm1-7 may be engaging A15 and U15 RNA in a different manner **(Fig. 1D)**. Specifically, PatMC/Lsm1-7 had a Hill coefficient of ∼1 for U15 RNA and ∼2 for A15 RNA, suggesting that binding to A15 is cooperative while U15 binding is not. This indicates that at least one or two copies of the PatMC/Lsm1-7 complex cooperatively bind to U15 or A15 RNA, respectively.

### Short RNAs are sufficient to promote dimerization of the PatMC/Lsm1-7 complex

To directly test the number of PatMC/Lsm1-7 complexes bound to short oligonucleotides, we used size exclusion chromatography coupled to multiangle light scattering (SEC-MALS). The SEC step fractionates protein complexes by hydrodynamic radius and molar mass is concurrently detected by light-scattering and differential refractometry (Wyatt 1993). The PatMC/Lsm1-7 complex was incubated with stoichiometric amounts of A15 RNA and subjected to SEC-MALS. The A15 RNA promoted the formation of two peaks that had identical protein composition and molar masses corresponding to that of a dimeric (two copies of PatMC/Lsm1-7) and tetrameric assembly **(Fig. 2A, Table 1)**. Shorter RNAs, such as A10, also promoted oligomeric PatMC/Lsm1-7 assemblies, with molar masses corresponding to dimeric and tetrameric complexes **(Fig. 2B)**. However, we observed that the A10 RNA reduced tetramerization and instead produced predominantly dimeric PatMC/Lsm1-7 complexes, based on the relative ratio of the peaks in the chromatogram. This suggests that RNA length many influence tetramerization, but short RNAs still promote higher order assembly of the PatMC/Lsm1-7 complex.

**Table 1:** Expected and observed molar masses from SEC-MALS. Expected and observed molar masses of SEC-MALS chromatograms. Error is displayed in parenthesis to the right of observed peak. Peak 1 and 2 refers to the earlier and later elution volume peaks, respectively.

| **Protein + RNA Complex** | **Expected Monomer Molar Mass (kDa)** | **Observed Molar Mass, Peak 1 (kDa)** | **Observed Molar Mass, Peak 2 (kDa)** |
| --- | --- | --- | --- |
| PatMC/Lsm1-7 | 133.5 | 304.5 (± 1.20%) | 160.3 (± 0.93%) |
| PatMC/Lsm1-7 + A15 | 138.2 | 576.0 (± 1.28%) | 273.3 (± 0.56%) |
| PatMC/Lsm1-7 + A10 | 136.6 | 554.2 (± 0.80%) | 275.5 (± 0.26%) |
| PatMC/Lsm1-7 + U15 | 137.8 | 548.1 (± 0.91%) | 268.1 (± 0.88%) |
| Lsm1-7 + U15 | 85.6 | 86.1 (± 2.40%) | NA |

**Figure 2:**
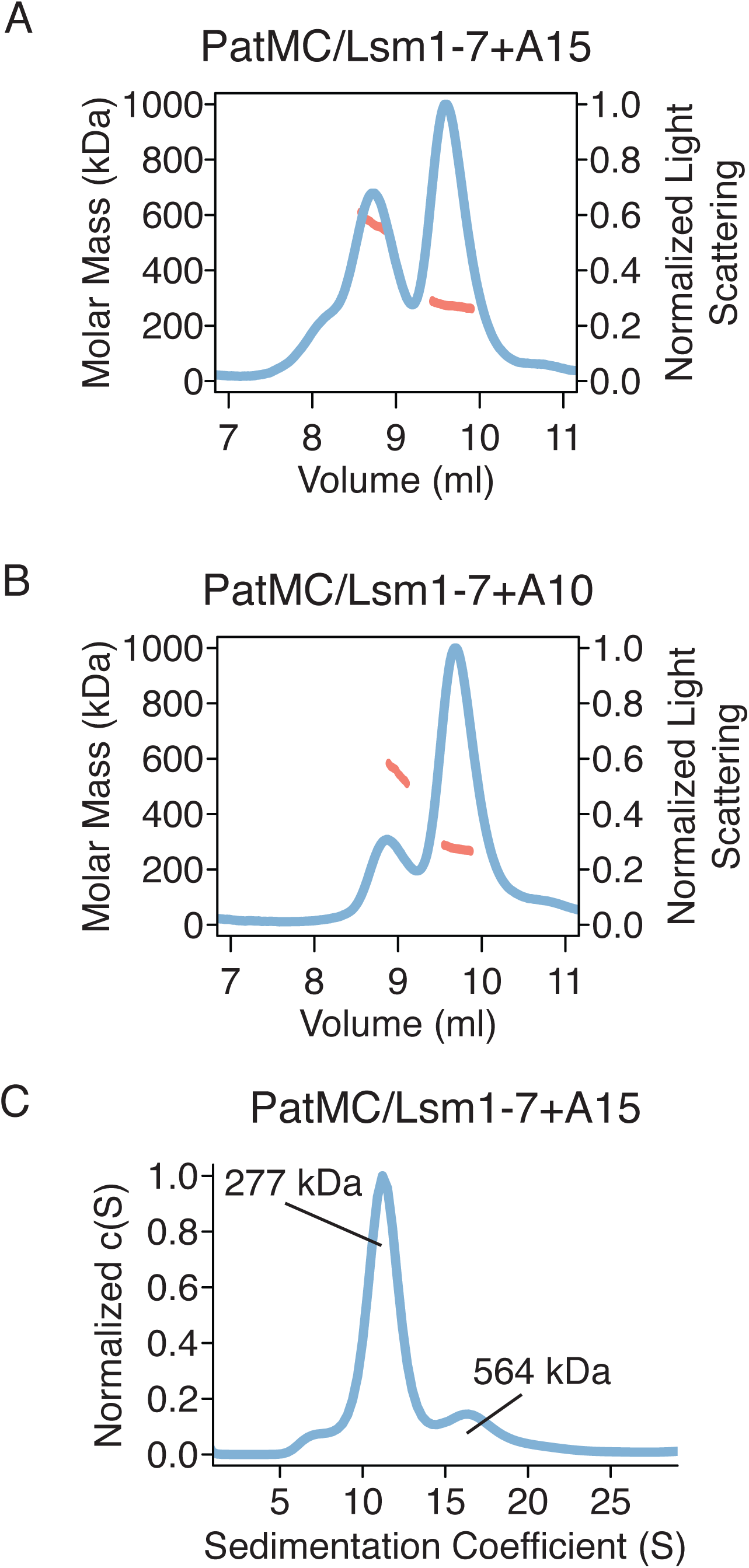
RNA promotes stable dimerization of the PatMC/Lsm1-7 complex. **A-B**, SEC-MALS of 20.6 *µ*M PatMC/Lsm1-7 with **A**, A15 and **B**, A10. Expected and observed molar masses are shown in Table 1. **C**, SV-AUC of 9.3 *µ*M PatMC/Lsm1-7 with stoichiometric amounts of A15 RNA at 250 mM NaCl. c(S) is the sedimentation distribution with molecular weights determined from fits.

To evaluate the stability of the oligomeric assemblies, we performed sedimentation velocity analytical ultracentrifugation (SV-AUC) on PatMC/Lsm1-7 with stoichiometric amounts of A15 RNA over the course of 12 hours. While we could detect a strong dimeric peak in the sedimentation distribution, there was minimal amounts of tetrameric assemblies **(Fig. 2C)**. Furthermore, the tetrameric fraction of the PatMC/Lsm1-7/A15 complex disassembled into dimers and tetramers upon reinjection over a size exclusion column, while the dimeric peak did not further dissociate (**Fig. S2A-C)**. This suggests that the tetrameric PatMC/Lsm1-7/RNA is less stable the dimeric species.

We next asked how PatMC/Lsm1-7 assembled on U15 RNA. As seen with A15 RNA, addition of stoichiometric amounts of U15 to the PatMC/Lsm1-7 complex resulted in two peaks by SEC-MALS with molar masses corresponding to dimeric and tetrameric assemblies **(Fig. 3A/B)**. This effect depends on PatMC, because Lsm1-7 alone bound to U15 RNA remained monomeric (**Fig. 3C**). Taken together, this indicates that PatMC promotes the higher order assembly of the PatMC/Lsm1-7 complex on both A15 and U15 RNA. This suggests that while both A15 and U15 promote dimerization of the PatMC/Lsm1-7 complex, some of the contacts of the dimer may differ, as evidenced by the difference in RNA binding cooperativity (**Fig. 1D**).

**Figure 3:**
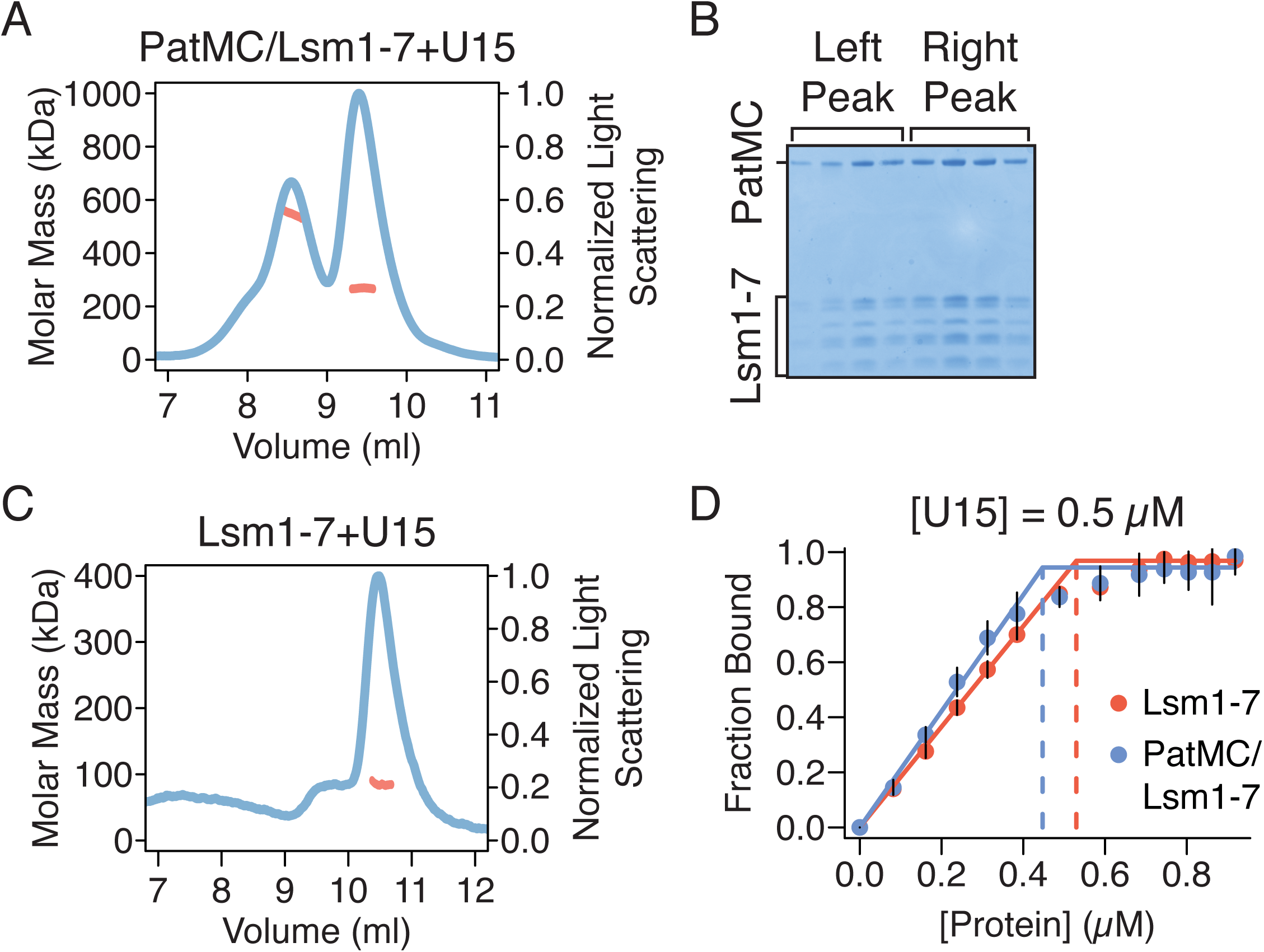
Multiple RNA sequences drive higher order PatMC/Lsm1-7 assembly in a Pat1 dependent manner. **A**, SEC-MALS of 20.6 *µ*M PatMC/Lsm1-7 with U15 RNA. The expected molar mass of the monomeric PatMC/Lsm1-7 complex is 133 kDa. **B**, Representative SDS-PAGE gel of fractions in **A. C**, SEC-MALS of 20.6 *µ*M Lsm1-7 with U15 RNA. The expected mass of the monomeric Lsm1-7 complex is 81 kDa. Expected and observed molar masses are shown in Table 1. **D**, Stoichiometry analysis of Lsm1-7 or PatMC/Lsm1-7 with 5’-FAM labelled U15 RNA at 0.5 *µ*M, which is >100-fold above K_d_.

It is possible that that each PatMC/Lsm1-7 complex binds an individual RNA or that multiple PatMC/Lsm1-7 complexes co-occupy a single RNA to promote oligomerization. To test these possibilities, we determined the stoichiometry of PatMC/Lsm1-7 binding to RNA. Experiments were performed under saturating conditions, where concentration of the oligo-RNA was far above the K_d_. Binding of labelled U15 was followed by fluorescence anisotropy. Titration of Lsm1-7 alone or the PatMC/Lsm1-7 complex results in saturation of the anisotropy signal at one equivalent of RNA, indicating 1:1 binding between PatMC/Lsm1-7 and the U15 oligonucleotide (**Fig. 3D**). Similar results were obtained for PatMC/Lsm1-7 binding to A15 (**Fig. S3**). This indicates that each PatMC/Lsm1-7 heterooctamer binds a single oligo-RNA, though we cannot exclude the possibility that PatMC/Lsm1-7 can co-occupy RNA sequences longer than 15nt tested here. Therefore, we conclude that RNA ligands of different sequences and lengths induce stable dimerization of the PatMC/Lsm1-7 complex.

### Dimerization is an intrinsic property of the PatMC/Lsm1-7 complex

As PatMC/Lsm1-7 binds to RNA as a higher order assembly, we asked if the complex had the intrinsic ability to multimerize in the absence of RNA. While PatMC/Lsm1-7 initially purified as a monomer, concentration and subsequent SEC-MALS of the complex in the absence of RNA revealed two peaks with molar masses corresponding to monomeric and dimeric PatMC/Lsm1-7 complexes **(Fig. 4A-C)**. This indicates that the PatMC/Lsm1-7 complex has the inherent ability to form multimers independent of nucleic acid, and suggests that RNA may drive higher order assembly.

**Figure 4:**
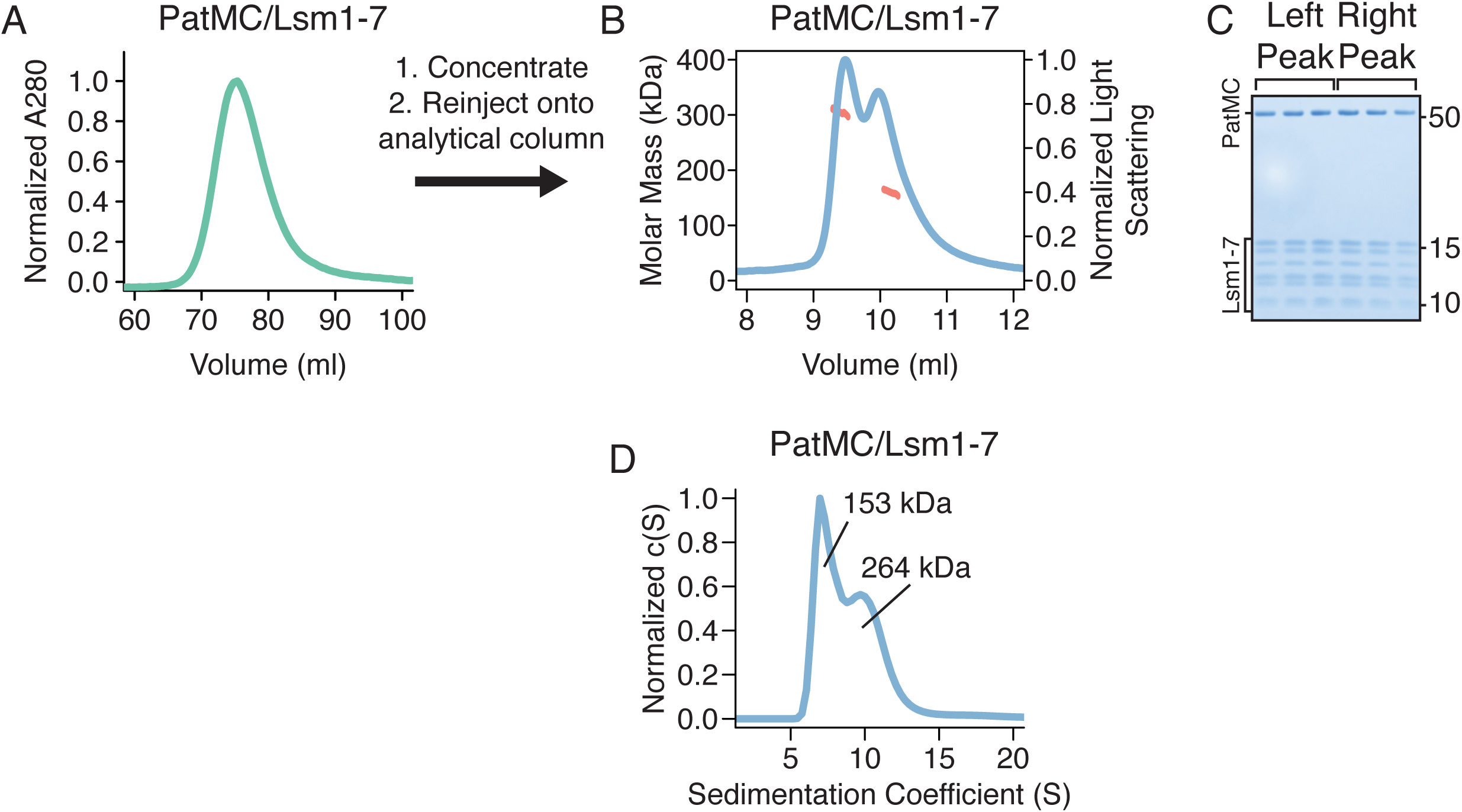
The PatMC/Lsm1-7 complex can intrinsically form a dimeric complex. **A**, Preparative size exclusion chromatography of the PatMC/Lsm1-7 complex in a 400 mM NaCl buffer. **B**, SEC-MALS of the concentrated PatMC/Lsm1-7 complex at 250 mM NaCl. The expected mass of the monomeric PatMC/Lsm1-7 complex is 133 kDa. Expected and observed molar masses are shown in Table 1. **C**, Representative SDS-PAGE gel of peaks (Left to right: earlier to later elution volumes). **D**, SV-AUC of 9.3 *µ*M PatMC/Lsm1-7 at 250 mM NaCl. c(S) is the sedimentation distribution with molecular weights determined from fits.

To test the stability of the assemblies in the absence of RNA, we performed SV-AUC on the PatMC/Lsm1-7 complex alone. Over the course of 12 hours, we observed both a monomeric and dimeric species, indicating that both these complexes were stable **(Fig. 4D)**. Furthermore, increasing salt concentrations favored monomerization of the complex, indicating PatMC/Lsm1-7 oligomerization is reversible **(Fig. S4)**. Because Lsm1-7 was monomeric in the absence of PatMC, the above results indicate PatMC drives multimerization of Lsm1-7, which may be enhanced by RNA binding (**Fig. 2A & 3A)**.

### PatMC promotes liquid-droplet formation with Dcp2 and recruits additional mRNA decay factors

Previous studies demonstrate that the monomeric, globular C-terminal domain of Pat1 can bind helical leucine motifs (HLMs) in the disordered C-terminus of Dcp2 (Charenton et al. 2017; Lobel et al. 2019). Known dimeric HLM binding proteins, such as Edc3, can interact with Dcp2 and undergo liquid-liquid phase separation with Dcp2 constructs that contain multiple HLMs (Schutz et al. 2017; Fromm et al. 2014). Our biochemical data demonstrate that PatMC can inherently oligomerize in the absence of RNA, so we tested if it could promote liquid-liquid phase separation with multivalent Dcp1/Dcp2 complexes, analogous to Edc3. PatMC was purified fused to maltose-binding protein (MBP) to enhance its solubility (see Methods). It was then mixed with a Dcp2 construct containing both the catalytic core and three HLMs in the disordered C-terminus extension, along with its obligate cofactor Dcp1 (Dcp1/Dcp2 1-504, termed Dcp1/Dcp2_Ext_) **(Fig. 5A)**. Though neither PatMC nor Dcp1/Dcp2_Ext_ formed condensates individually, mixing stoichiometric amounts of Dcp1/Dcp2_Ext_ with MBP-PatMC resulted in formation of phase separated droplets (**Fig. 5B and data not shown)**.

**Figure 5:**
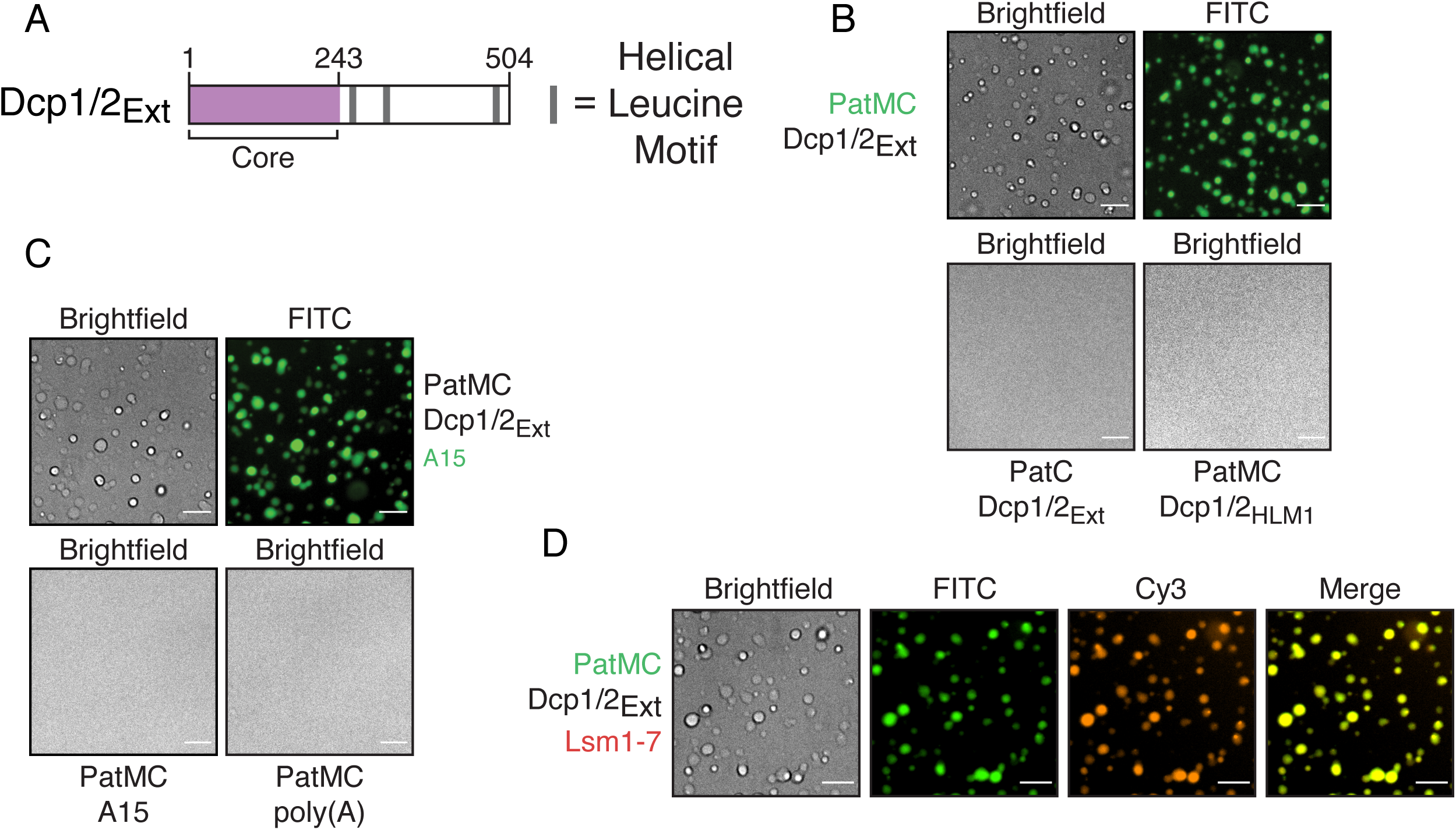
Oligomerization of PatMC promotes liquid-liquid phase separation with Dcp2 and recruitment of additional RNA decay machinery. **A**, Schematic of Dcp2 construct used. Purple represents the globular domain that cleaves the m7G cap. Gray bars represent helical leucine motifs in the disordered C-terminal extension. **B**, Brightfield and fluorescence microscopy of droplets with 2.5 *µ*M Pat constructs (0.1 *µ*M FITC-labeled) with stoichiometric amounts of Dcp1/Dcp2_Ext_. **C**, Brightfield and fluorescence microscopy of droplets with 2.5 *µ*M PatMC and 2.5 *µ*M rA15 RNA or 13.4 ng/*µ*l poly(A) RNA with or without 2.5 *µ*M Dcp1/Dcp2_Ext_ and 0.1 *µ*M FAM-rA15. **D**, Brightfield and fluorescence microscopy of 2.5 *µ*M PatMC (0.1 *µ*M FITC-labelled) and Dcp1/Dcp2_Ext_ with 0.1 *µ*M Alexa555 labelled Lsm1-7. All images taken at 40x magnification. Scale bar, 10 *µ*m.

To understand the requirements of PatMC and Dcp1/Dcp2_Ext_ for droplet formation, we queried how individual regions of both complexes contribute to phase separation. The C-terminal domain of Pat1 is monomeric and binds HLMs, but did not promote phase separation of Dcp1/Dcp2_Ext_ (**Fig. 5B**). Additionally, PatMC did not phase separate with a Dcp1/Dcp2 complex containing a single HLM (Dcp2 residues 1-266, termed Dcp2_HLM1_) **(Fig. 5B)**. These data suggest that the middle and C-terminal domains of Pat1 promote phase separation of Dcp1/Dcp2 by driving self-association and binding HLMs on Dcp2, respectively. We conclude PatMC oligomerization promotes phase separation with multivalent cofactors such as Dcp2.

PatMC and Dcp2 both interact with RNA, which can trigger or potentiate liquid-liquid phase separation with oligomeric RNA binding proteins (RBPs) (Mugler et al. 2016; Lin et al. 2015). The PatMC/Dcp1/Dcp2_Ext_ droplets were able to incorporate A15 RNA, though A15 RNA did not change the critical concentration required for phase separation (**Fig. 5C & S5A)**. However, neither short A15 RNAs nor poly(A) RNA promoted droplet formation with either PatMC or Dcp1/Dcp2_Ext_ alone (**Fig. 5C and data not shown**). RNA could not trigger phase separation with either Dcp1/Dcp2_Ext_ or PatMC alone, suggesting that the protein-protein interactions are the primary driver of phase separation between PatMC and Dcp2.

We next asked if other mRNA decay factors could be incorporated in PatMC/Dcp1/Dcp2 droplets. Lsm1-7, but not a nonspecific protein, was recruited of pre-formed PatMC/Dcp1/Dcp2_Ext_ condensates, indicating that PatMC can bridge both 5’ (Dcp1/Dcp2) and 3’ (Lsm1-7) decay factors in the context of the phase separated droplet (**Fig. 5D & S5C**). Lsm1-7 neither phase separates with PatMC nor affected the critical concentration for droplet formation, consistent with Lsm1-7 being a monomeric complex (**Fig. S5A & S5B**). Furthermore, Dcp1/Dcp2_Ext_/PatMC/Lsm1-7 condensates recruited RNA (**Fig. 5SD**). These observations suggest that Pat1 may bridge both 5’ and 3’ activities in the context of a phase separated droplet.

## DISCUSSION

Our biochemical reconstitution uncovers how Pat1 broadens the specificity of the Lsm1-7 complex and promotes higher order assembly of multiple mRNA decay factors. First, Pat1 expands the Lsm1-7 complex’s sequence preference by enhancing binding to adenine-rich RNAs. Second, PatMC promotes cooperative binding of Lsm1-7 to RNA, which drives oligomerization on nucleic acid. Third, the PatMC/Lsm1-7 complex has the inherent ability to oligomerize, which is dependent on Pat1 and consistent with coimmunoprecipitation data in fission yeast (Wang et al. 2017). Finally, we show that an oligomeric PatMC drives phase separation with multivalent Dcp1/Dcp2 complexes that can recruit RNA and additional decay factors to droplets. Taken together, this biochemically reconstituted system reveals how Pat1 increases the range of RNA targets bound by the Lsm1-7 complex and facilitates higher order assembly of multiple decapping factors (**Fig. 6A/B)**.

**Figure 6:**
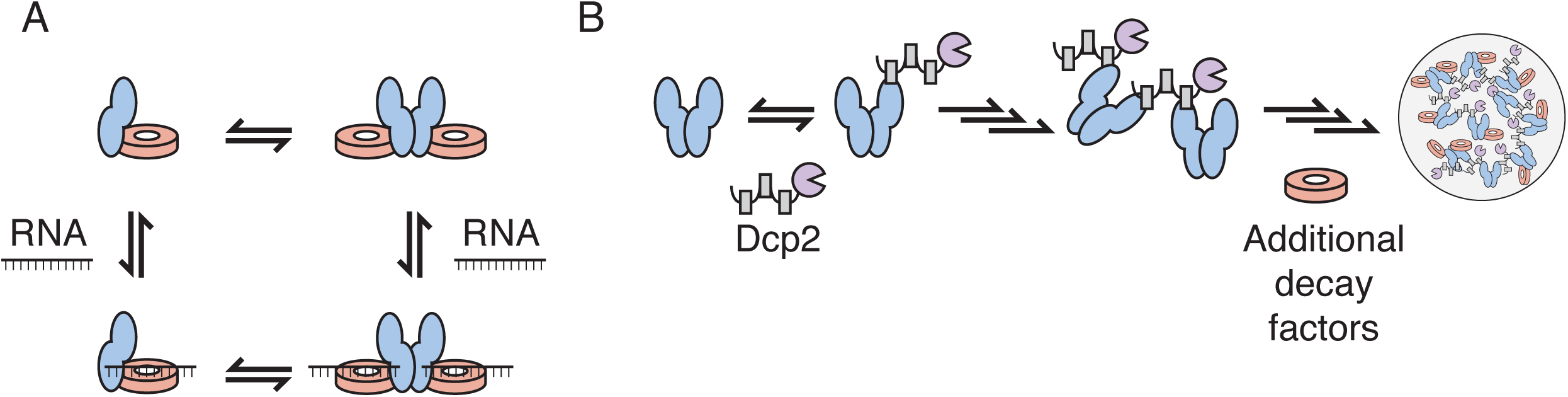
Model for how Pat1 increases specificity and assembly for different mRNA decay factors. **A**, Proposed thermodynamic coupling between Pat1 (blue), Lsm1-7 (red), and RNA binding to promote multimerization of the Pat1/Lsm1-7 complex. **B**, Binding of oligomeric Pat1 assemblies to multivalent Dcp1/Dcp2 complexes promotes phase separation and recruitment of additional mRNA decay factors.

The Lsm1-7 ring is the high affinity RNA binding component of the Pat1/Lsm1-7 complex and has a preference for U-rich oligonucleotides (Wu et al. 2014; Lobel et al. 2019; Chowdhury et al. 2007). Pat1 broadens the specificity of the Lsm1-7 complex by enhancing the affinity of Lsm1-7 for oligonucleotides with higher adenine content (Lobel et al. 2019; Chowdhury et al. 2013) (**Fig. 1**). Because PatMC does not bind RNA with appreciable affinity on its own, we suggest that Pat1 may allow the Lsm1-7 complex to bind sequences which it has inherently weaker affinity for and therefore expand the complex’s target repertoire (Lobel et al. 2019) (**Fig. S1G**).

Though PatMC increases the range of RNA substrates bound by Lsm1-7, it is unclear if all RNA targets bind in the same manner. For example, PatMC does not enhance the affinity of Lsm1-7 for U15 RNA, in contrast to A15 RNA or A/U rich RNA. Moreover, higher adenine contents favor more cooperative binding of PatMC/Lsm1-7. On the other hand, Pat1/Lsm1-7 stoichiometrically binds all RNA targets as a stable dimer. While we observe tetramers with 15mer RNAs, these assemblies are less stable than the dimeric species (**Fig. 2 & Fig. S2)** Determining the binding modes of different RNAs with the Pat1/Lsm1-7 complex remains a challenge for future structural studies.

PatMC also consistently increased the cooperativity of Lsm1-7 binding to all oligonucleotides tested, suggesting a coupling between protein-protein interactions and RNA binding. In support, addition of RNAs drive formation of stable dimeric PatMC/Lsm1-7 assembly whereas in the absence of RNA, PatMC/Lsm1-7 exists in monomer/dimer equilibrium. This suggests two possible pathways which the PatMC/Lsm1-7 complex can load onto RNA. First, RNA may bind to dimeric PatMC/Lsm1-7 from a pre-existing monomer/dimer equilibrium. Alternatively, RNA could bind monomeric PatMC/Lsm1-7 which then forms a dimer (**Fig. 6A**). These pathways, could in fact, be part of a thermodynamic cycle, which is in qualitative agreement with our observations.

The inherent ability of PatMC to oligomerize drives phase separation with multivalent cofactors, such as Dcp2 containing multiple HLMs. This derives from multivalent interactions between Dcp2 and oligomeric PatMC. These droplets can recruit Lsm1-7, providing evidence that Pat1 can bring both 5’ and 3’ mRNA decay factors in close proximity in the context of these phase separated droplets (**Fig. 5**). Additional partners of Pat1, such as Dhh1, may cooperate to further promote droplet formation (Sachdev et al. 2019). Nucleating high local concentration of multiple decapping RBPs in the context of a phase separated droplet may be leveraged for 5’ and 3’ end communication during decay (**Fig. 6B**). Future work is required to understand how an oligomeric Pat1 is regulated and functions in assembling an active decapping mRNP during 5’-3’ mRNA decay.

The discovery that Pat1 has the ability to oligomerize is reminiscent of the homo-hexameric bacterial Lsm-family protein, Hfq, and its cofactor Crc. Recent work has demonstrated that two copies of Crc can bridge two Hfq hexamers in an RNA dependent manner (Pei et al. 2019; Sonnleitner et al. 2018). While the details of higher order Lsm assemblies between bacteria and eukaryotes differ, oligomerization may be a conserved feature of Lsm complexes and their cofactors.

## MATERIALS AND METHODS

### Protein expression and purification

All proteins were expressed in BL21(DE3)* (Thermofisher) cells in LB media. Cells were grown to OD_600_ = 0.6 at 37 °C, after which IPTG was added to 1 mM. Cells were then grown overnight at 18 °C for 16 hours. For expression of copurified PatMC/Lsm1-7, a polycistron containing all seven Lsm proteins was cloned into site one of a pET-Duet vector, with an N-terminal hexahistidine tag followed by a TEV cleavage sequence on Lsm1. A codon optimized PatMC (residues 296-754) was ordered from IDT and cloned into site two of the pET-Duet vector. Cells were harvested by centrifugation and lysed in appropriate buffer. For Lsm1-7 and PatMC/Lsm1-7 complexes, cells were lysed in Buffer A (2 M NaCl, 20 mM HEPES pH 7.5, 20 mM Imidazole, 5 mM βME, protease inhibitor (Roche)) by sonication. Lysate was subsequently clarified by centrifugation and the supernatant was bound to Ni-NTA resin (GE) at 4 °C for 1 hour. The resin was then transferred to a gravity column and washed with 20 column volumes of Buffer A before being eluted in 25 mL of Buffer E (250 mM NaCl, 250 mM Imidazole, 20 mM HEPES pH 7.0, 10 mM βME). The elution was then loaded directly onto a 5 mL HiTrap Heparin column (GE). The heparin column was run at 2 ml/min from a 0.25-1 M NaCl gradient over 20 column volumes. Fractions containing the appropriate protein complex were concentrated in 30 kD concentrators (Amicon) to ∼2 mL before adding TEV overnight at 4 °C. The following day, the sample was filtered and further purified by gel filtration using a Superdex 200 16/60 column (GE). Coexpressed PatMC/Lsm1-7 was purified in 400 mM NaCl, 20 mM HEPES 7.0, 1 mM DTT and Lsm1-7 alone was purified in 150 mM NaCl, 20 mM HEPES pH 7.0, 1 mM DTT. Fractions containing protein were concentrated before being flash frozen and stored at −80 °C.

All MBP-Pat1 fusions were purified as described previously (Lobel et al. 2019). For Spycatcher purification, a C-terminal KCK tag was added (SpycatcherKCK) and purified similar to the MBP-Pat1 fusions, with the heparin step omitted. SpycatcherKCK was purified on Superdex 75 16/60 column in 150 mM NaCl, 20 mM HEPES pH 7.0, 0.5 mM TCEP.

### Fluorescence polarization

All fluorescent polarization experiments were performed in 200 mM NaCl, 20 mM HEPES pH 7.0, 1 mM DTT, 5 mM MgCl_2_ with 0.3 *µ*g/ul Acetylated BSA (Promega). All RNAs used were labelled with 5’ FAM (IDT) and were used at final concentration of 500 pM. All binding curves were fit to the following Hill model for single site binding:

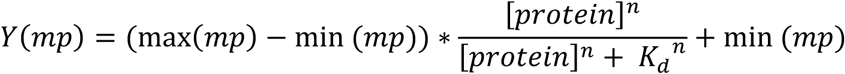

To determine the ΔG of binding, each independent replicate was fitted to the above model and ΔG was determined by the relationship ΔG = +RT*ln(K_d_). The ΔG from each individual fit was averaged and plotted with standard deviation. Hill coefficients were averaged from fitting three separate binding isotherms and shown with standard deviation.

For analysis of stoichiometry, 5’-FAM RNA was kept at 0.5 *µ*M in the same buffer as used for the fluorescence polarization assay. Protein was titrated into solution containing labelled RNA and fluorescence polarization was measured for each concentration. The linear portion was fit to a linear model and the average of the last four points were used to fit a line at the saturation point. The intersection of these two lines was used to determine the binding stoichiometry.

### Analytical Size Exclusion Chromatography (SEC) and SEC-MALS

Analytical SEC was performed in buffer M (250 mM NaCl, 20 mM HEPES pH 7.0, 1 mM DTT), or in the appropriate NaCl concentration. Samples were mixed at ∼30 *µ*M and incubated for 15 minutes in 350 mM NaCl, 20 mM HEPES pH 7.0, 1 mM DTT on ice before being filtered and injected onto a GE Superdex 200 10/300-Increase analytical size exclusion column. When appropriate, samples were mixed with 1.1-fold molar excess RNA. All samples were run at 0.35 ml/min, and peaks were analyzed by SDS-PAGE (Invitrogen). For experiments involving reinjection of fractions over SEC, 500 *µ*l of fractions were spin filtered before reinjecting over SEC.

For SEC-MALS, 165 *µ*g of sample was filtered through a 0.1 *µ*m spin filter (Amicon) before being injected onto a pre-equilibrated KW-804 column (Shodex, New York, New York). For samples with RNA, stoichiometric amounts of RNA were added prior to spin filtration. Data was acquired with an inline DAWN HELEOS MALS and Optilab rEX differential refractive index detector (Wyatt Technology, Santa Barbara, CA). All analysis was performed using ASTRA VI software (Wyatt Technology). Data was then exported and plotted with R.

### Analytical ultracentrifugation

Sample was buffer exchanged into buffer M using Zeba spin columns (Thermofisher) and diluted to 9.3 *µ*M. When appropriate, stoichiometric amounts of RNA were added to the sample after buffer exchange. AUC cells were assembled according to manufacturer’s protocol and 100 *µ*l of sample was loaded into the cell. The sample was incubated at 22 °C for >2 hours prior to centrifugation. Samples were run at 30,000 rpm for 10-12 hours in a Beckman XL/A analytical ultracentrifuge. Scans for samples containing only protein were collected at 280 nm, and samples containing RNA were scanned at both 280 and 260 nm. Sedimentation velocity analysis was performed in SEDFIT (NIH) and plots were generated with GUSSI (Schuck 2003; Brautigam 2015). Experimental parameters were determined using SEDNTERP (NIH). The following parameters were used for fitting: partial volume, 0.739818; buffer density, 1.0101; buffer viscosity, 0.0104032.

### Protein labelling

For labelling with dyes, proteins were buffer exchanged into appropriate labelling buffer using Zeba spin columns (Thermofisher). Lsm1-7 and SpycatcherKCK were labelled with 5-fold molar excess Alexa Fluor 555 maleimide for one hour at room temperature in 150 mM NaCl, 20 mM HEPES pH 7.5, 0.5 mM TCEP. Reactions were quenched by addition of βME to a final concentration of 10 mM. MBP-PatMC was labelled with 4-fold molar excess NHS-Fluorescein (Thermofisher) for 1 hour at room temperature in 150 mM NaCl, 150 mM Sodium bicarbonate pH 8.4 before being quenched by adding TRIS-HCl pH 8.0 to a final concentration of 50 mM. All quenching steps were performed at room temperature for 20 minutes. Free dye was separated from labelled protein by Illustra NICK columns (GE) according to the manufacturer’s instruction. Labelling efficiency and concentrations were calculated by UV-vis spectroscopy.

### Microscopy

All images were acquired with Nikon Eclipse T*i* equipped with a 40x dry lens. Samples were prepared in a 384 well plate (Greiner) that was cleaned with 0.1M NaOH and passivated with PEG-silane and 100 mg/ml BSA (Sigma-Aldrich) before being washing with water to remove residual BSA. Proteins or RNA were mixed at specified concentrations in a final buffer concentration of 60 mM NaCl, 20 mM HEPES pH 7.0, 1 mM DTT. When appropriate, dye-labelled protein or RNA were added to 100 nM. Samples were incubated at room temperature for 20 minutes prior to imaging. Images were analyzed in FIJI (Schindelin et al. 2019).

## ACKNOWLEDGEMENTS

We thank Alexandra Rizo and Serena Sanulli for experimental guidance, the Nikon Imaging Center at UCSF for use of the microscope, Daniel Southworth’s lab for use of SEC-MALS, and Ryan Tibble and Nathan Gamarra for comments on the manuscript. This work was supported by US National Institutes of Health grant R01GM078360 to J.D.G.

## SUPPLEMENTAL FIGURE LEGENDS

**Supplemental Figure 1:**
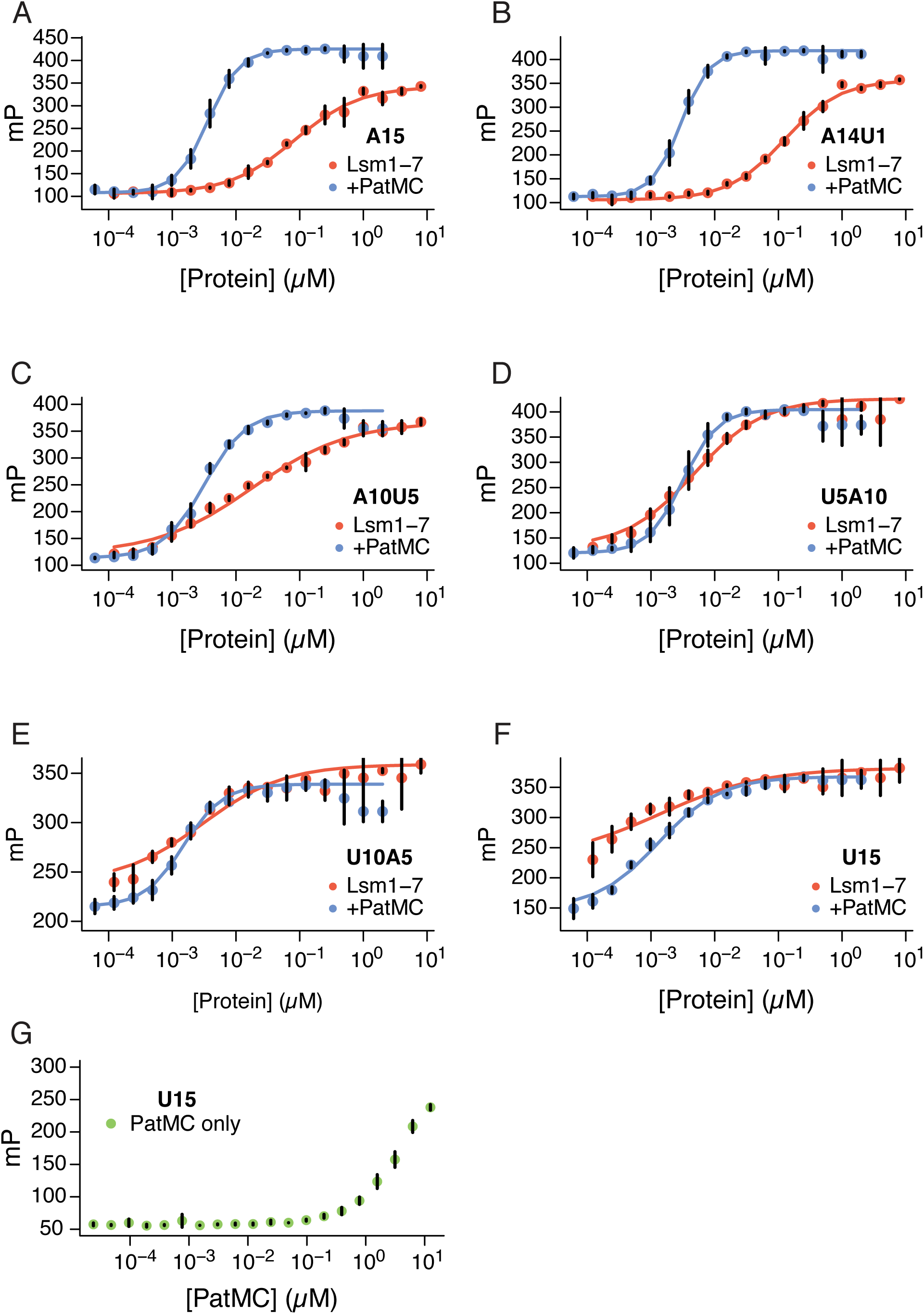
Binding of Lsm1-7 +/- PatMC to different RNAs. **A-F**, Lsm1-7 or PatMC/Lsm1-7 binding to **A**, A15 **B**, A14U1 **C**, A10U5 **D**, U5A10 **E**, A5U10 **F**, U15 (n = 3 for all). **G**, MBP-PatMC binding to U15 in 150 mM NaCl (n = 2).

**Supplemental Figure 2:**
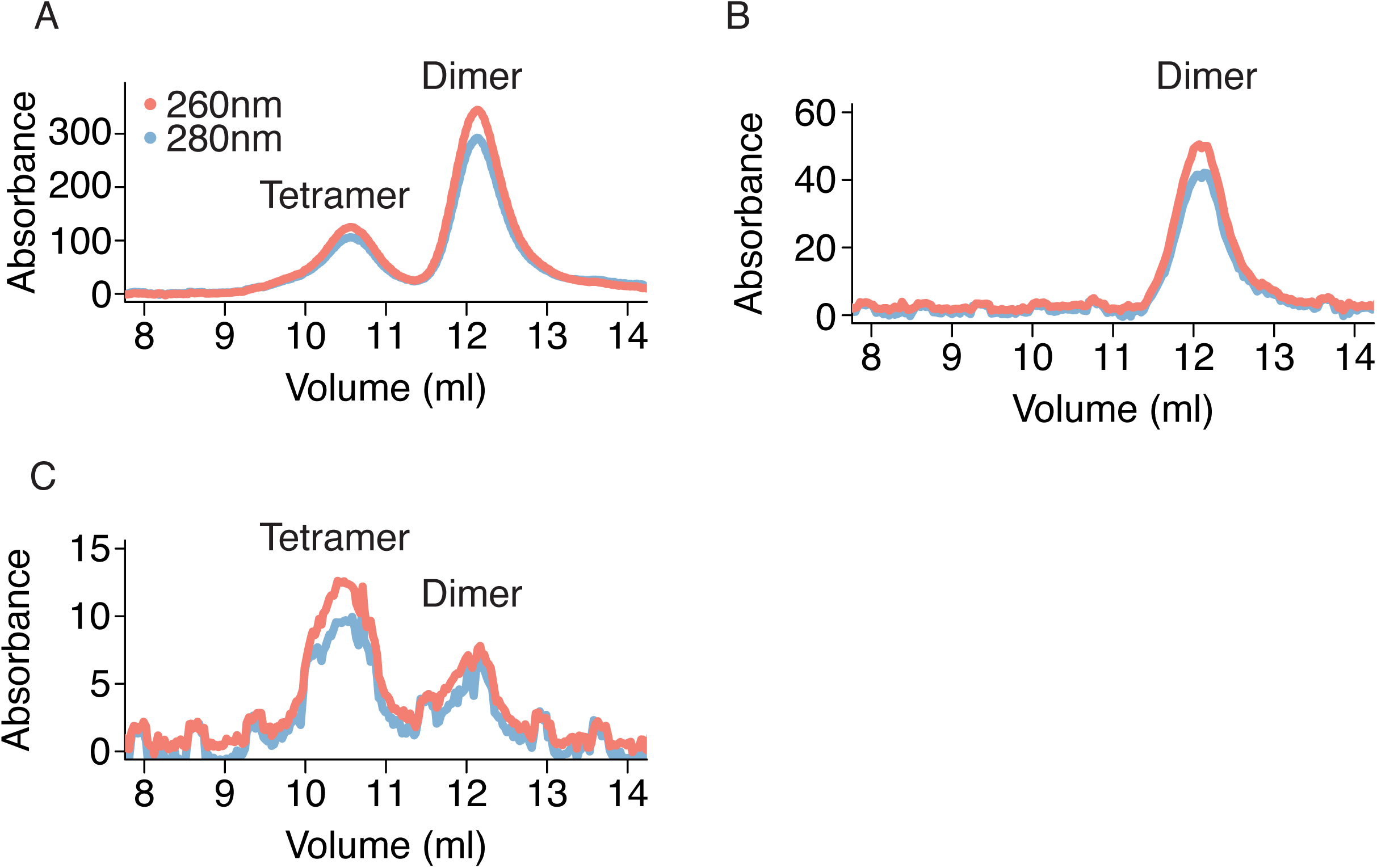
The dimeric PatMC/Lsm1-7/RNA assembly is stable. **A**, Analytical size exclusion chromatography of PatMC/Lsm1-7 with A15 RNA. Peaks corresponding to dimer and tetramer are labelled. **B**, Reinjection of peak corresponding to the dimeric PatMC/Lsm1-7 complex. **C**, Reinjection of the peak corresponding to the tetrameric PatMC/Lsm1-7 complex. All conditions were in a 250 mM NaCl buffer. The 260 and 280 nm absorbances are displayed in red and blue, respectively.

**Supplemental Figure 3:**
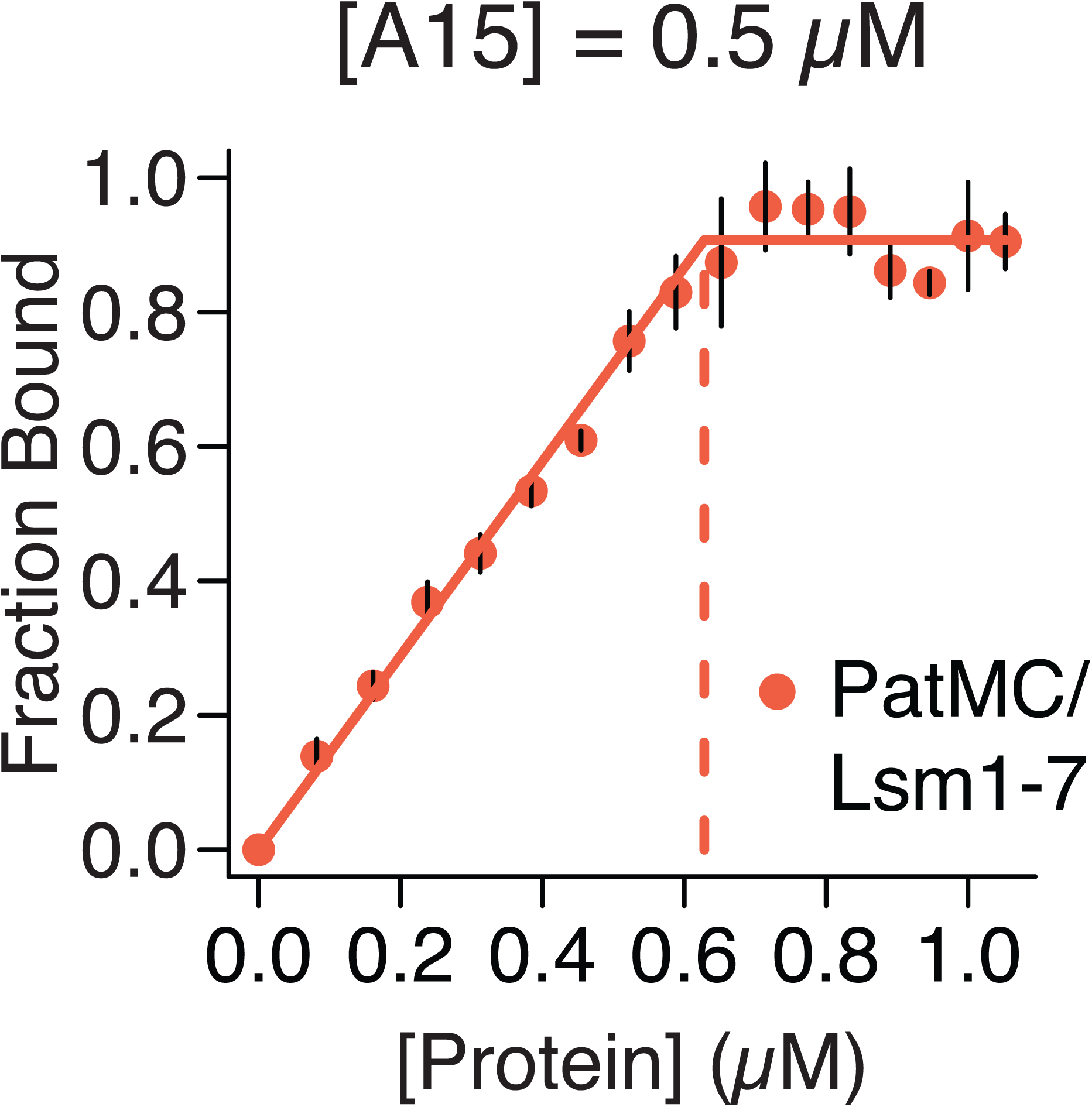
PatMC/Lsm1-7 binds A15 stoichiometrically. Stoichiometry analysis of Lsm1-7 or PatMC/Lsm1-7 with 5’-FAM labelled A15 RNA at 0.5 *µ*M, which is ∼100-fold above K_d_.

**Supplemental Figure 4:**
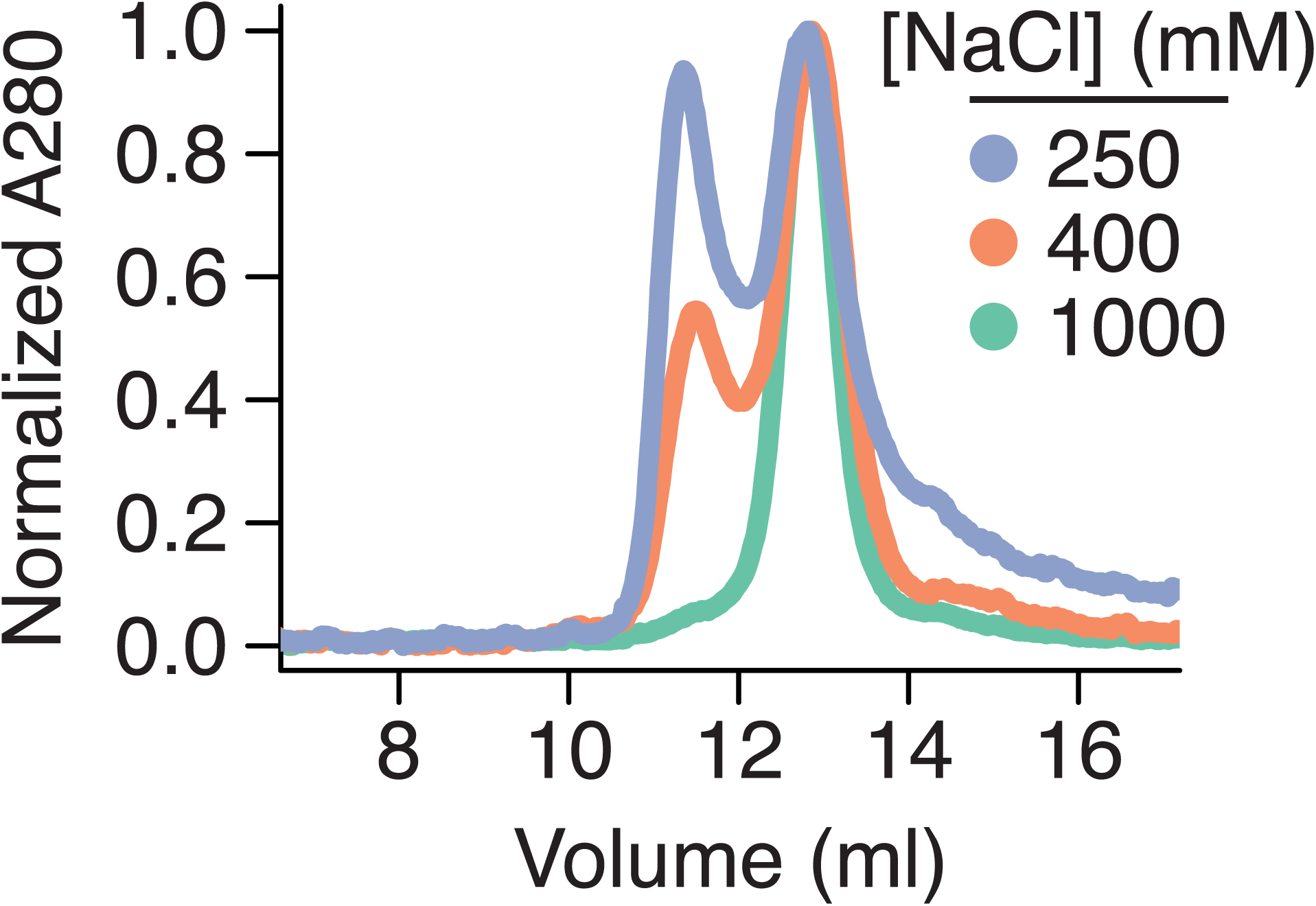
The monomer/dimer equilibrium of the PatMC/Lsm1-7 complex is sensitive to ionic strength of solution. Analytical size exclusion chromatography of PatMC/Lsm1-7 complexes in different ionic strength solutions as indicated in the figure. All protein was run in 20 mM HEPES pH 7.0, 1 mM DTT and the specified NaCl concentration.

**Supplemental Figure 5:**
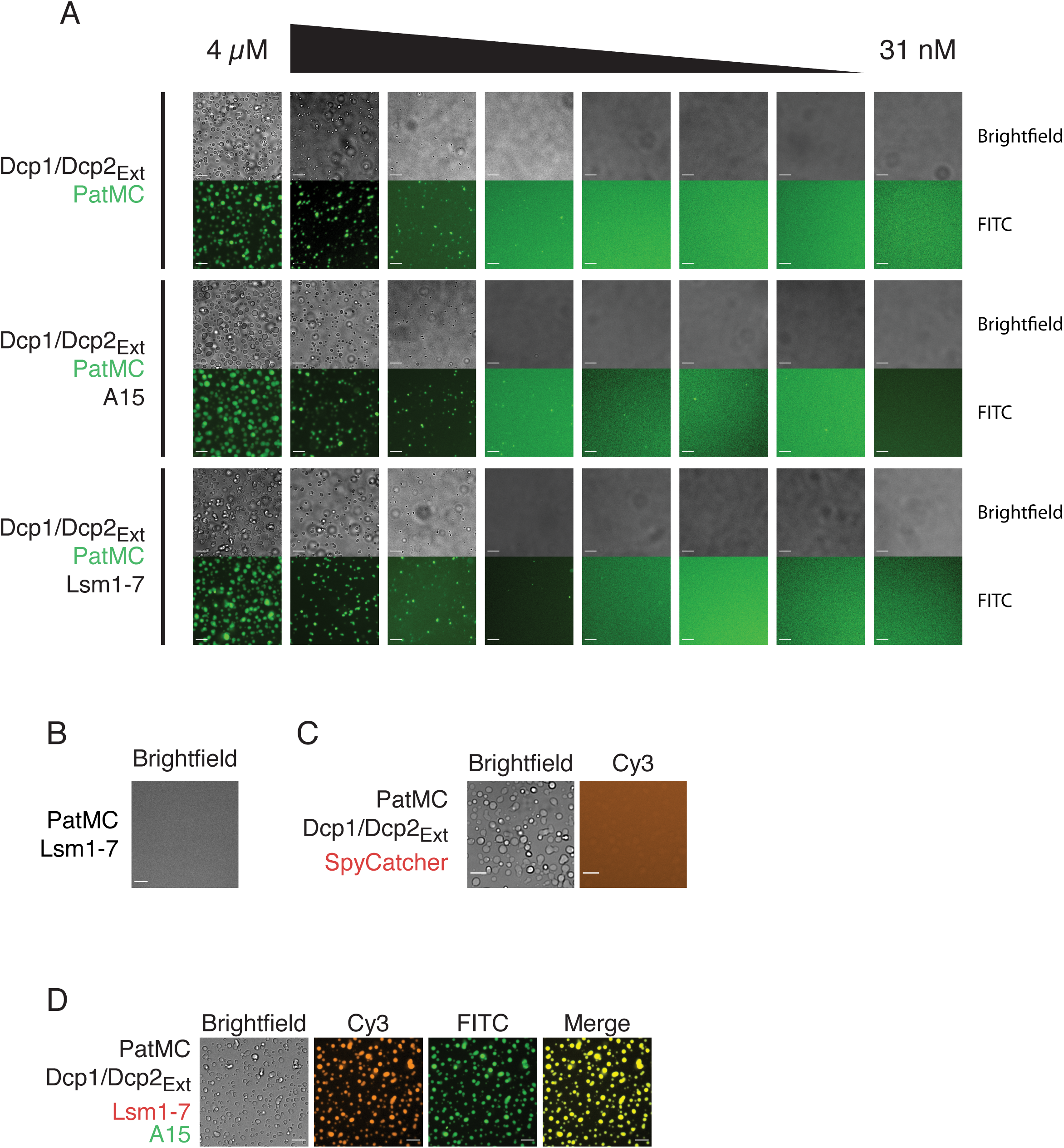
The multivalent PatMC/Dcp2_Ext_ interaction is required for liquid-liquid phase separation. **A**, Brightfield images of serial dilutions of droplets containing MBP-PatMC/Dcp1/Dcp2_Ext_ with A15 RNA or Lsm1-7. (0.1 *µ*M MBP-PatMC labelled). **B**, Brightfield images of 2.5 *µ*M PatMC and Lsm1-7 **C**, Brightfield and fluorescence microscopy of 2.5 *µ*M MBP-PatMC and Dcp1/Dcp2_Ext_ with 0.1 *µ*M Alexa555-Spycatcher labelled. **D**, Brightfield and fluorescence microscopy of 2.5 *µ*M MBP-PatMC/Dcp1/Dcp2_Ext_/Lsm1-7 droplets with 0.1 *µ*M FAM-rA15 and 0.1 *µ*M Alexa555-Lsm1-7. All images taken at 40x magnification. Scale bar, 10 *µ*m.

